# Integrated reanalysis of global riverine fish eDNA datasets shows robustness and congruence of biodiversity conclusions

**DOI:** 10.1101/2025.09.16.676481

**Authors:** Yan Zhang, Heng Zhang, Hiroshi Akashi, Camille P. Albouy, Kara J. Andres, José Barquín, Jeanine Brantschen, Richard E. Connon, Joseph M. Craine, Deirdre Gleeson, Alejandra Goldenberg-Vilar, Alexia M. González-Ferreras, Chelsea Hatzenbuhler, Kamil Hupało, Josephine Hyde, Wataru Iwasaki, Mark D. Johnson, Aron D. Katz, Vyacheslav V. Kuzovlev, Courtney E. Larson, Laurène A. Lecaudey, Florian Leese, Matthieu Leray, Feilong Li, Till-Hendrik Macher, Quentin Mauvisseau, María Morán-Luis, Georgia Nester, Helio Quintero, Tsilavina Ravelomanana, Merin Reji Chacko, Mattia Saccò, Naiara Sales, Tamara Schenekar, Martin Schletterer, Saskia Schmidt, Nicholas O. Schulte, Robin Schütz, Jinelle H. Sperry, Emma R. Stevens, Sarah A. Stinson, Steven Weiss, Fei Xia, Hui Zhang, Song Zhang, Wenjun Zhong, Shuo Zong, Loïc Pellissier, Xiaowei Zhang, Florian Altermatt

## Abstract

The analysis of environmental DNA (eDNA) has revolutionized biodiversity assessments in aquatic ecosystems, enabling non-invasive monitoring of fish communities across diverse regions. However, the global comparability of these eDNA datasets remains ambiguous due heterogeneous sampling protocols and bioinformatic workflows across studies, particularly regarding the robustness of their conclusions on biodiversity assessments. Here, we conducted a meta-analysis of 58 riverine fish eDNA metabarcoding datasets, covering 1,818 sampling sites worldwide, to evaluate the robustness of eDNA-derived biodiversity patterns. We found that species richness estimates and metrics of community structure derived under a common bioinformatic workflow were overall consistent with those of original analyses, despite the relatively high variability in bioinformatic analyses in the respective original studies. Contrastingly, congruence of species identity varied more extensively across datasets, mostly reflecting different completeness and regional relevance of reference databases. Restricting taxonomic assignment to basin-specific species pools improved species identification accuracy, while datasets lacking publicly accessible or well-curated reference data were more prone to mismatches. Year of sampling had a positive effect on taxonomic congruence, such that more recent studies showed increased robustness, also reflecting improved reference database coverage and enhanced species-level identification over time and overall method congruence in more recent years. Overall, the suitability and potential of eDNA for global biodiversity monitoring is corroborating overall robust biodiversity estimates, irrespective of the bioinformatic approaches. Our study underlines the effectiveness and need of further harmonization of bioinformatic workflows and strengthened region-specific reference databases for improved taxonomic resolution and comparability across studies.

## Introduction

Freshwater ecosystems, particularly riverine systems, are important biodiversity hotspots, but face some of the highest rates of biodiversity loss (Almond et al., 2022; Hughes et al., 2021b). Halting this decline is a central target in both the Global Sustainable Development Agenda and the Kunming-Montreal Global Biodiversity Framework (CBD, 2022; Obrecht et al., 2021; Tickner et al., 2020). Effective and sustainable monitoring of river biodiversity is a fundamental step in achieving these global biodiversity goals (Gonzalez et al., 2023). However, significant gaps in biodiversity assessment persist, particularly in the Global South, where freshwater biodiversity is often rich yet poorly studied (Chapman et al., 2024). These gaps hinder effective conservation strategies and policy development, leaving many regions and species underrepresented or unidentified (Chapman et al., 2024; Senior et al., 2024), and there is an urgent need for rapid, cost-effective, and standardized tools to assess biodiversity across riverine ecosystems worldwide.

Environmental DNA (eDNA) has emerged as a transformative tool for biodiversity monitoring, offering a non-invasive method to detect species across ecosystems, with particular effectiveness in riverine ecosystems (Altermatt et al., 2025; Blackman et al., 2024; Deiner et al., 2017). By capturing and analyzing genetic material from environmental samples, eDNA enables species identification without the need for direct organismal observation (Pawlowski et al., 2020; Taberlet et al., 2012). The approach is particularly valuable for studying rare and cryptic species, such as fish, which are often difficult to sample using conventional gillnet or electrofishing techniques (Cilleros et al., 2019; Yao et al., 2022). Additionally, eDNA facilitates the integration of genetic signals across river networks, providing a more comprehensive understanding of biodiversity by accounting for species’ distribution across large spatial scales (Carraro et al., 2020; Urycki et al., 2024). The ability to detect rare and cryptic species further enhances the potential of eDNA, allowing for the survey of understudied species that might otherwise remain undetected by traditional monitoring methods (Piggott et al., 2021).

While eDNA holds great promise for global biodiversity assessments (Altermatt et al., 2025), achieving comparability across existing datasets remains a significant challenge (Blackman et al., 2024; Loeza-Quintana et al., 2020). Several aspects of sample collection and processing, including primer selection (Yang et al., 2023; Zhang et al., 2020), sample volume (Sakata et al., 2020; Zhang et al., 2023), sampling strategy (Zhang et al., 2023) and sequencing platforms (Korostin et al., 2020; Singer et al., 2019), have been shown to influence species detection and biodiversity estimates. However, less attention has been given to subsequent bioinformatics workflows and bioinformatic analysis of the data (but see Marques et al. (2020) and Mathon et al. (2021)). The specifics of bioinformatic workflow and analysis may not only significantly affect species identification and the consistency of results across studies (Mathieu et al., 2020), but also hold enormous potential for consistent (re)analyses under common bioinformatic pipelines and procedures. It is this long-term value of eDNA metabarcoding results, namely that the sequences can be reanalyzed and used across studies, that gives the method a high applicability for global (meta)analyses and intercalibration and use of data. To effectively achieve comparability and robustness across studies, an understanding of the consistency and variability introduced by bioinformatic approaches and reference databases used is critical. Variations in bioinformatic pipelines, including sequence quality filtering, clustering methods, and reference database quality, can lead to discrepancies in species identification (Keck et al., 2022b; Marques et al., 2020), such that meaningful comparisons between studies are not a priori guaranteed. Specifically, data cleaning steps, including raw read quality control, clustering criteria, and abundance/occurrence thresholds, can impact the biodiversity estimates (Marques et al., 2020). The stringency of quality control affects the rates of false negatives, particularly for rare species that fall below detection thresholds, while less rigorous filtering can allow contaminant ASVs/OTUs to remain, leading to false positives that distort biodiversity estimates (Zinger et al., 2019). Further, the choice and quality/extent of reference database used for matching sequences to taxonomic units is particularly important and profoundly affect species identification (Keck et al., 2022b; Weigand et al., 2019). Standardized bioinformatics workflows will be essential for improving the reliability and comparability of eDNA-based biodiversity assessments, and will enable a new, integrative view on biodiversity globally.

Here, we addressed comparability and integration of eDNA studies by conducting a global meta-analysis of eDNA datasets focused on riverine fish communities, integrating a broad set of published and unpublished eDNA datasets. Specifically, we reanalyzed 58 original raw sequencing datasets using common bioinformatic pipelines, addressing three key questions: (1) How congruent are fish biodiversity patterns, focusing on both the species richness and community structure, when reanalyzing them in one common bioinformatic approach and comparing to the original individual analyses? (2) To what extent do species identities align across eDNA datasets? (3) How much does the outcome of these comparisons depend on methodological choices? We hypothesized that the congruence between reanalyzed and original eDNA data is influenced by several factors, and particularly focused on effects of geographic context such as the ecoregion where the study was conducted, methodological choices including the barcode or primer used and sequencing depth, as well as the completeness and type of reference database used in the original taxonomic assignment. The latter includes whether a public global or custom-built reference database was used, and the year of sampling, approximating the level of reference database completeness at the time the study was conducted and indicating method improvements over time in a rapidly developing field. By addressing these questions and the respective influence of these factors, we aim to improve our understanding of how eDNA-based biodiversity assessments can be standardized and integrated and to provide recommendations for enhancing the comparability and reliability of eDNA studies in general.

## Method

### Datasets compilation

A comprehensive literature review was conducted on December 13, 2022, using the Web of Science Core Collection, with the full search query provided in the Supplementary Text. The initial search returned 673 records, which were first filtered by removing 13 book chapters, leaving 660 records. These records were sorted by relevance and manually reviewed for inclusion based on the following criteria: 1) literature type—review and meta-analysis papers were excluded; 2) target taxonomic group—only studies focused on fish communities were considered; 3) ecosystem— only studies on rivers or streams were included; 4) biomonitoring method—only studies using eDNA metabarcoding were kept.

After this initial screening, 50 relevant papers were selected. These papers were further assessed for data availability, with the following criteria for inclusion: 1) studies that provided geographic coordinates for sampling sites; 2) studies that reported species richness or provided raw sequencing data. This second review led to the retention of 42 papers. From these, metadata were extracted, including publication details, data availability, sampling protocols, metabarcoding procedures, and bioinformatic methods (Supplementary Table 1).

To expand our dataset and capture additional relevant studies, we made a public call on social media platform X (back then known as Twitter), targeting researchers and institutions involved in similar studies. This outreach aimed to gather unpublished or less accessible data to enhance the comprehensiveness of our datasets. As a result, 23 datasets were provided by co-authors, contributing data from 588 sampling sites across 19 river catchments. Of the provided datasets, one was from Africa, five from Asia, two from Australia, seven from Europe, five from North America, and three from South America. Three of the datasets had already been included during the paper review and screening process, bringing the total to 62 datasets (see “DataID” in Supplementary Table 1). In Supplementary Table 1, metadata on all of these datasets were provided, covering details on sampling (e.g., time, frequency, number of sites, volume), metabarcoding experiments (e.g., DNA extraction, PCR, sequencing), and bioinformatics (e.g., quality control, clustering, taxonomy assignment) aspects. For each dataset, raw sequencing data, species richness estimates for each sampling site (species richness table), and species-site matrixes were extracted. Four studies, each with only a single sampling site, were excluded, resulting in a final set of 58 datasets used in the meta-analysis (Figure 1).

**Figure 1.**
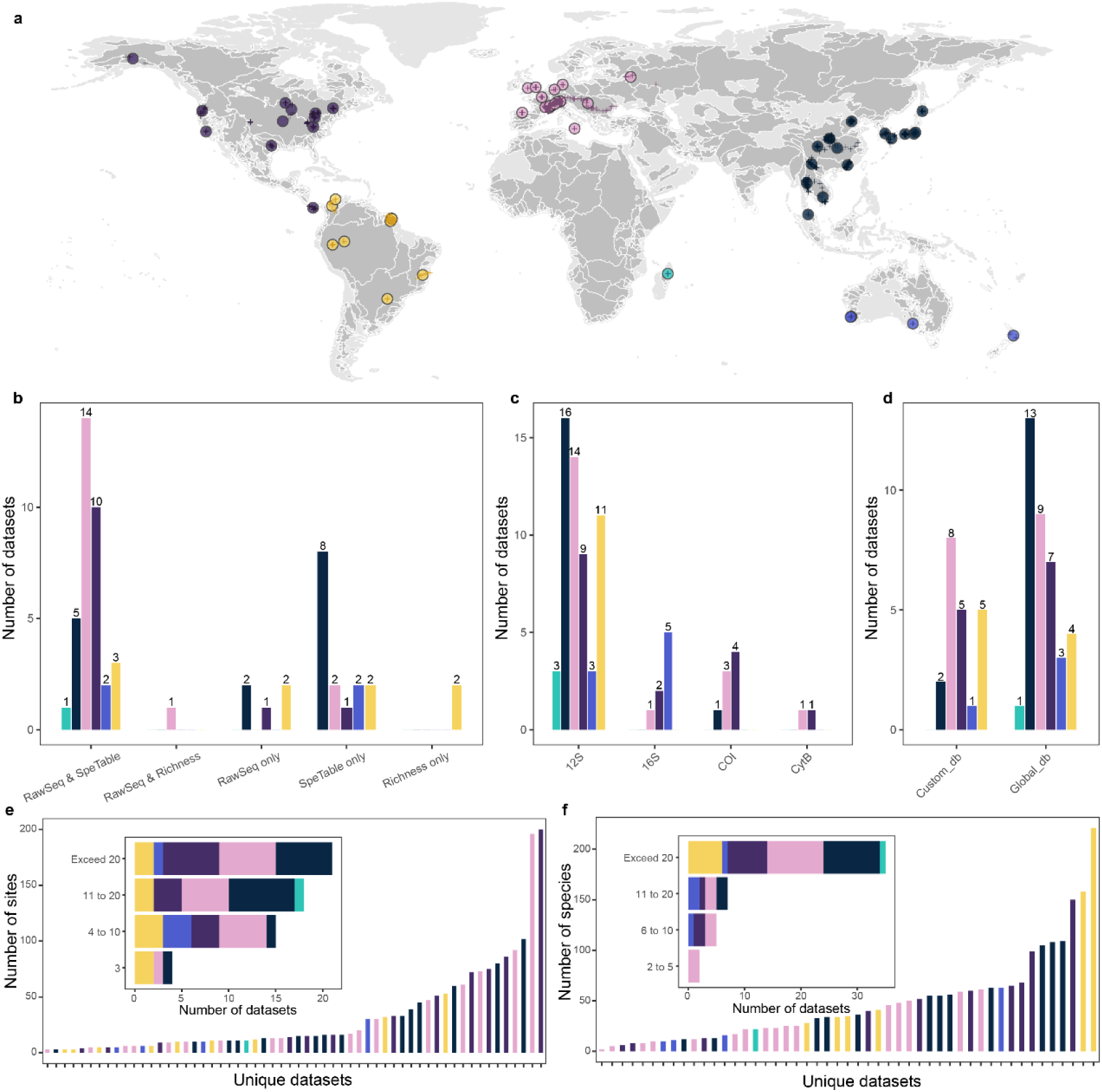
Distribution and summary of the 58 datasets. Panel (a) shows the geographical distribution of the 58 datasets (circles) and 1,818 sampling sites (crosses). Light gray areas represent all global river basins, while dark gray areas indicate the river basins covered by the 58 datasets. Panels (b–d) summarize the available data types (raw sequencing data, species-site table, and species richness), barcode region (12S, 16S, COI, or CytB), and the type of reference database used in the original species assignments (global or custom). Panel (e) summarizes the number of sites included in the datasets. Panel (f) provides a summary of the number of fish species in the originally classified datasets. The colors used in all panels correspond to the six ecoregions.

Among them, 41 datasets contained raw sequencing data, with some using multiple primers (Figure 1). To account for this, the datasets were further separated based on unique dataset-primer combinations, resulting in a total of 56 distinct raw sequencing datasets, each archived in separate folders (see “RawSeqID” in Supplementary Table 1). Primer names were standardized based on their sequences, and sample names were checked and standardized to align with the species-site matrixes or species richness tables. All fish species names were standardized using “rfishbase” R package (Froese & Pauly, 2002).

### Reference database curation

The CRUX pipeline, the first module of the Anacapa Toolkit (Curd et al., 2019), was used to curate reference databases for specific primer sets. Briefly, the process begins with in-silico PCR using ecoPCR on the EMBL nucleotide sequence database to generate a seed library of taxon-specific identifiers (Ficetola et al., 2010; Stoesser et al., 2002). The pipeline then verifies the amplicon match by checking primer regions and trimming them using CUTADAPT (Martin, 2011). Since many GenBank sequences do not include primer regions, CRUX performs two rounds of BLAST against the NCBI nucleotide database (Camacho et al., 2009). The first-round filters for full-length reads, while the second includes shorter reads (down to 70% of the full length). The results are de-replicated, retaining only the longest reads. CRUX also generates a taxonomy file using entrez-qiime, mapping reads to taxonomic categories (Baker, 2016). Reads with incomplete or unclear taxonomy (e.g., “unassigned”, “uncultured” are excluded to maintain high-quality reference data. The parameters for each primer set are outlined in Supplementary Table 2. Finally, the reference databases are formatted into Bowtie2-formatted index libraries.

### Bioinformatics Workflow

For raw sequencing datasets that were not demultiplexed, paired-end reads were first merged using VSEARCH (Rognes et al., 2016). The *barcode_splitter* was used for demultiplexing raw sequencing reads based on barcode sequences, allowing up to three mismatches (Leach & Parsons, 2019). Any reads that did not match a barcode or had multiple barcode matches were discarded. Demultiplexed reads were renamed and organized into separate fastq files for each sample. Additionally, *NGSFILTER* from the OBITools package was used for filtering and processing in cases double-barcode adapters were used (Boyer et al., 2016).

The DADA2 pipeline was then run in the R environment to process the demultiplexed sequencing reads (Callahan et al., 2016). Initially, reads were imported and filtered to remove ambiguous bases using the *filterAndTrim()* function, with a maximum of 0 ambiguous bases. Primer sequences were identified and removed using CUTADAPT, with primer sequences checked in all orientations to ensure accurate trimming (Martin, 2011). Low-quality reads were discarded using the *filterAndTrim()* function, with a maximum expected error rate (maxEE) of 2 and a truncation quality (truncQ) threshold of 2. Error rates were learned using *learnErrors()*, and reads were denoised with the *dada()* function to generate amplicon sequence variants (ASVs). Paired-end reads were merged using *mergePairs()*, and chimeric sequences were removed using *removeBimeraDenovo()*. A sequence table was created, retaining only chimera-free ASVs. The number of reads at each step of processing (raw, filtered, denoised, merged, and non-chimeric) was tracked for each sample.

The taxonomy of the ASVs was assigned using the Classifier module of the Anacapa Toolkit, which employs *Bowtie2* and a Bayesian Least Common Ancestor (BLCA) algorithm (Curd et al., 2019; Gao et al., 2017; Langmead & Salzberg, 2012). Initially, all ASVs were globally aligned to the CRUX database using Bowtie 2. The top 100 hits were then processed using the BLCA script for taxonomic assignment. The confidence of each taxonomic assignment was assessed using bootstrap scores. The species annotations in the output were initially uncorrected based on the local species and were therefore referred to as “global assignments” throughout the manuscript.

### Local species record and reference database coverage

We used a global database of freshwater fish occurrence, compiled through an extensive literature review and online resources, which provides complete species lists, including well-known native, introduced and invasive fishes, for 3,119 drainage basins (Tedesco et al., 2017). This database was used to define the local species for each dataset, identified at three hierarchical scales: basin, country, and ecoregion. First, each dataset was assigned to a specific drainage basin. For datasets not directly linked to a basin in the database, the nearest available basin was used as a proxy. Fish records for the assigned basin, country, or ecoregion were then retrieved to compile the local species list for each dataset.

The barcode species coverage of local fish species was defined as the proportion of local species in each dataset represented in the barcode reference database (generated in section Reference database curation). We also compared the global assignments with the local species. These comparisons were made at the three geographical scales (basin, country, and ecoregion), using species or genus as the taxonomic resolution. To ensure consistency, the fish species names in the global assignments, barcode reference database, and local species lists were validated and standardized using the “rfishbase” R package (Froese & Pauly, 2002). The differences of the barcode reference database coverage of local species and the number of species assigned to local taxa in the global assignments between ecoregions were assessed by Kruskal-Wallis rank sum test (Ostertagova et al., 2014) followed by post-hoc Dunn’s test after checking the normality and homogeneity of variances (Supplementary Table 3). The *multcompLetters2()* function from the “multcompView” R package was used to extract group-wise significance letters based on *p* values from Dunn’s test.

To validate the local species lists and determine the appropriate geographical scale and taxonomic resolution for subsequent analyses, we re-examined the comparison between the global assignments and the local species lists for datasets from Switzerland (Europe), French Guiana (South America), the St. Regis River (North America), and the Chao Phraya River (Thailand), where historical fish records are publicly available (Supplementary Table 4).

### Comparison of species richness estimates and community structures

We compared species richness estimates between the original and reanalyzed assignments using linear mixed models (LME), with “DataID” as random factor (Pinheiro et al., 2013). For the reanalyzed assignments, we considered four types: a) global assignments (uncorrected using local fish lists), b) assignments corrected to the basin-scale local species, c) assignments corrected to the country-scale local species, and d) assignments corrected to the ecoregion-scale local species.

We conducted these comparisons for all datasets, as well as separately for datasets based on either a global or custom reference database used in the original assignments. The global reference database included sources such as GenBank, ENA, MitoFish. The custom reference databases were defined as: 1) using barcodes from local species sampled within the region, 2) curating a reference database from global reference database by retaining only barcodes of historically recorded species, and 3) assigning species against global reference database and then correcting based on local historical records.

To further assess the robustness of community structures across ecoregions and assignment types, we extracted the full species list for each dataset and each assignment type (original assignment, re-assignments at the global level, and re-assignments at the basin level), and combined these into a single species occurrence matrix. We then calculated the Jaccard distance and performed Non-metric Multidimensional Scaling (NMDS), followed by PERMANOVA and ANOSIM tests using “vegan” R package to evaluate the differences in community composition (Dixon, 2003).

### Comparison of species identity

The species identities were compared between the original assignments and the reanalyzed assignments corrected to the basin-level local species. All species were classified into four categories: 1) shared species, 2) species unique to the re-assignments, 3) species unique to the original assignments but considered local species at the basin level, and 4) species unique to the original assignments but not local species at the basin level. The proportions of shared species in the re-assignments and original assignments, as well as the number of additional local species found in the original assignments, were assessed for normality using the Shapiro-Wilk test and for homogeneity of variance using Levene’s test. The results indicated that the data violated assumptions of normality and homogeneity of variance (Supplementary Table 3), so instead of ANOVA, comparisons were made between ecoregions using Kruskal-Wallis rank sum test followed by post-hoc Dunn’s test (Ostertagova et al., 2014). The differences between datasets using either the global or custom reference database were assessed by the Wilcoxon test (Fix & Hodges Jr, 1955).

Generalized linear models (GLMs) were used to examine factors influencing the proportion of shared species between original and reanalyzed assignments, based on both species count and sequencing read abundance (Hastie & Pregibon, 2017). Predictor variables included median sequencing depth, year of sampling, reference database type (global vs. custom), and reanalyzed species richness. To compare relative abundance patterns, LME models were applied, comparing species unique to the reanalysis with those shared with the original assignments, with dataset identity included as a random effect (Pinheiro et al., 2013).

## Results

### Summary of the datasets

A total of 58 datasets (see “DataID” in Supplementary Table 1), encompassing 1,818 sampling sites globally, were included in the meta-analysis (Figure 1). These datasets were distributed across different ecoregions, with 15 from Asia, 17 from Europe, 12 from North America, 9 from South America, 4 from Oceania, and 1 from Africa. Among these, 11 datasets utilized multiple barcode markers. The majority (51 of 58) employed the 12S rRNA gene, followed by 6 using 16S rRNA, 7 using COI (Cytochrome C Oxidase subunit I gene), and 2 using CytB (Mitochondrially Encoded Cytochrome B gene). Regarding species assignment, 21 datasets used custom reference databases, while the remaining datasets relied on global reference databases, which did not account for historical species occurrences. The number of sampling sites per dataset ranged from 3 to 200, with 19 datasets containing fewer than 10 sites, 18 covering 11 to 20 sites, and 21 including more than 20 sites. The number of species detected per dataset ranged from 2 to 221, with 35 datasets identifying more than 20 species.

Of the 41 datasets with available raw sequencing data (Figure 1), seven employed more than one primer set. After organizing the raw sequencing data by dataset identity and primer used, we identified a total of 56 distinct raw sequencing datasets (see “RawSeqID” in Supplementary Table 1). Six of these lacked sequencing indexes required to distinguish individual samples, so finally 50 raw sequence datasets were reanalyzed using a standardized bioinformatics workflow.

### Reference database coverage of the local species

Out of the 50 raw sequencing datasets, 23 distinct primer sets were used, and reference databases were curated separately for each primer set. These primer sets covered a broad spectrum of species according to the in-silico results, with varying amplification capabilities against GenBank (Supplementary Table 5). Approximately 4,000 species had 12S rRNA barcodes corresponding to 11 primer sets, amplifying between 3,785 and 4,387 species—except for the Riaz02 primer set, which could only amplify 4 species. Five primer sets targeting 16S rRNA genes successfully amplified between 4,166 and 9,022 fish species, while five COI primer sets amplified between 5,307 and 9,761 species. In contrast, the CytB primer sets were more limited, amplifying 2,571 species with Cytb01 and 1,705 species with Cytb02.

The reference database coverage of local species was assessed across datasets at different geographical scales (basin scale, country scale, or ecoregion scale) and taxonomic resolutions (species and genus levels), revealing substantial variability based on both the barcode used and the ecoregion (Africa, Asia, Europe, North America, Oceania, or South America) of the datasets. At the species level, reference database coverage ranged from 8.2% to 80.9% at the basin scale, 9.2% to 76.4% at the country scale, and 8.8% to 61.6% at the ecoregion scale, excluding the poorly performing Riaz01 primer set. At genus level, the coverages were higher, ranging from 49.0% to 100% at the basin scale, 47.2% to 95.9% at the country scale, and 50.6% to 94.9% at the ecoregion scale. Comparisons of barcode coverage by different barcode regions showed significant differences at the species level at both the basin (Kruskall-Wallis H = 17.75 *p* = 0.001) and ecoregion (Kruskall-Wallis H = 20.23 *p* < 0.001) scales, as well as at the genus level at the ecoregion scale (Supplementary Figure 1, Kruskall-Wallis H = 21.57 *p* < 0.001). The datasets using 12S rRNA barcode generally exhibited a wide range of coverage for local species, with consistent patterns across geographical scales and taxonomic resolutions (Supplementary Figure 1, Supplementary Figure 2). Additionally, when comparing datasets from different ecoregions, reference database coverage of local species varied significantly across all geographical scales and taxonomic resolutions (Figure 2). Datasets from Asia, Europe, and North America exhibited similar coverage, with only minor variations, whereas datasets from Africa and South America showed insufficient coverage.

**Figure 2.**
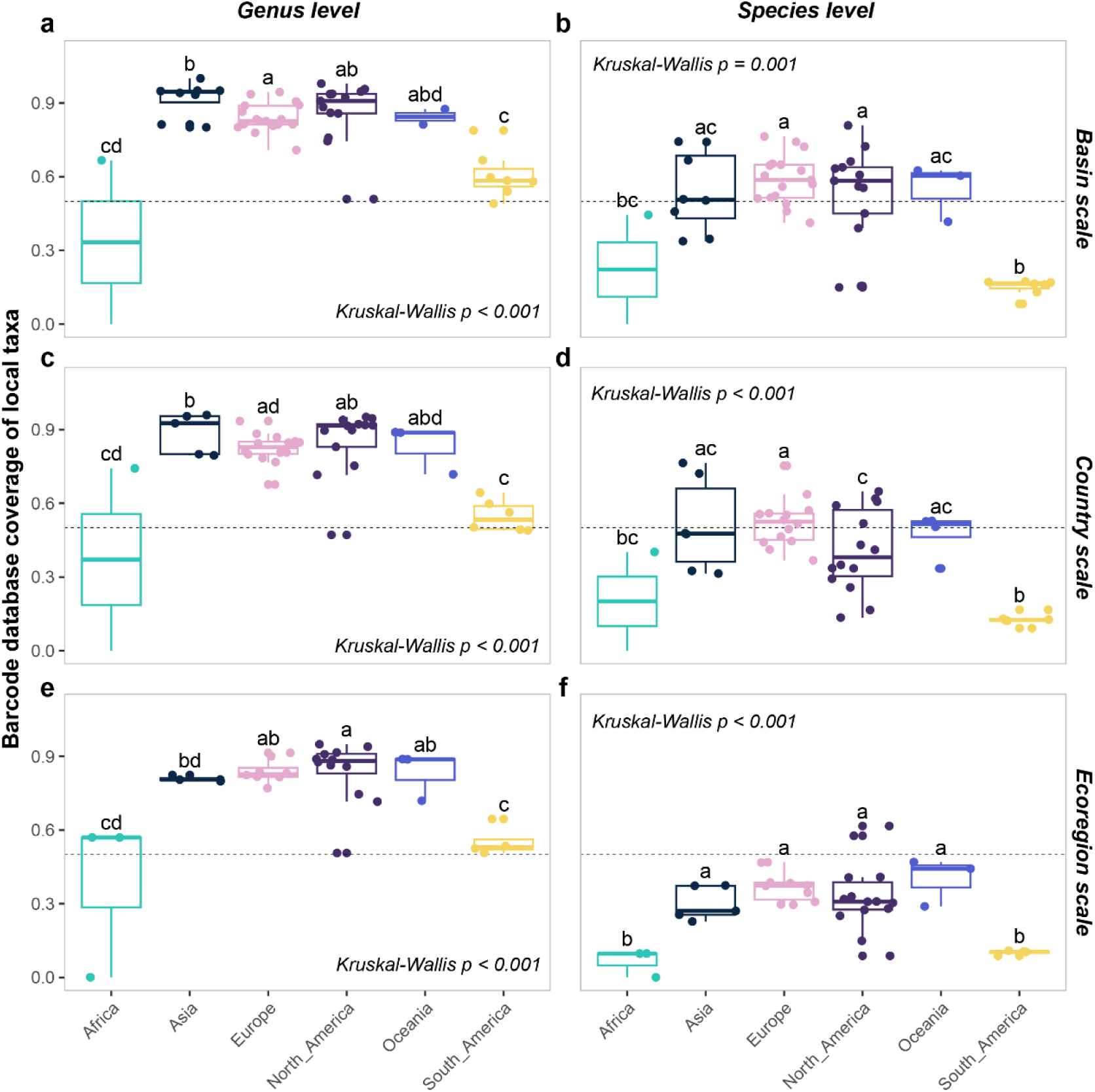
The barcode reference database coverage of local species across different spatial scales for each dataset. Local species were defined at the basin (a–b), country (c–d), and ecoregion (e–f) scales. Panels (a), (c), and (e) illustrate the genus-level coverage, while panels (b), (d), and (f) show the species-level coverage. The differences between ecoregions were assessed by ANOVA and subsequent pairwise Dunn’s tests. Different letters indicate statistically significant differences between groups (p < 0.05). Groups sharing the same letter are not significantly different, whereas groups with no letters in common differ significantly. The horizontal dotted lines indicate a value of 0.5.

The global assignments were compared with local species lists at different geographical scales, revealing significant differences between datasets from various ecoregions (Figure 3 and Supplementary Figure 3). While there were no significant differences in the proportion of species assigned as local species/genus between different barcodes, datasets from North America showed the highest proportion of local species. At the basin scale, an average of 63.3% of species and 81.5% of genera were assigned as local. Over 80% of species could be assigned as local at the country and ecoregion scales. In contrast, datasets from Asia and Europe showed lower local species assignment at the basin scale (30.1% for Asia and 36.2% for Europe), with higher proportions at the ecoregion scale (53.4% for Asia and 49.6% for Europe). For Oceania, Africa, and South America, local species assignment was notably lower. In Oceania, an average of 9.5% of species and 17.6% of genera were assigned as local at the basin scale, with coverage increasing to 53.3% of species at the country scale. In Africa, 11.2% of species and 13.8% of genera were assigned as local at the basin scale, rising to 27.9% of species at the country scale, and 49.9% of species at the ecoregion scale. South America showed 19.8% of species assigned as local at the basin scale, with 29.1% at the country scale and 54.8% at the ecoregion scale.

**Figure 3.**
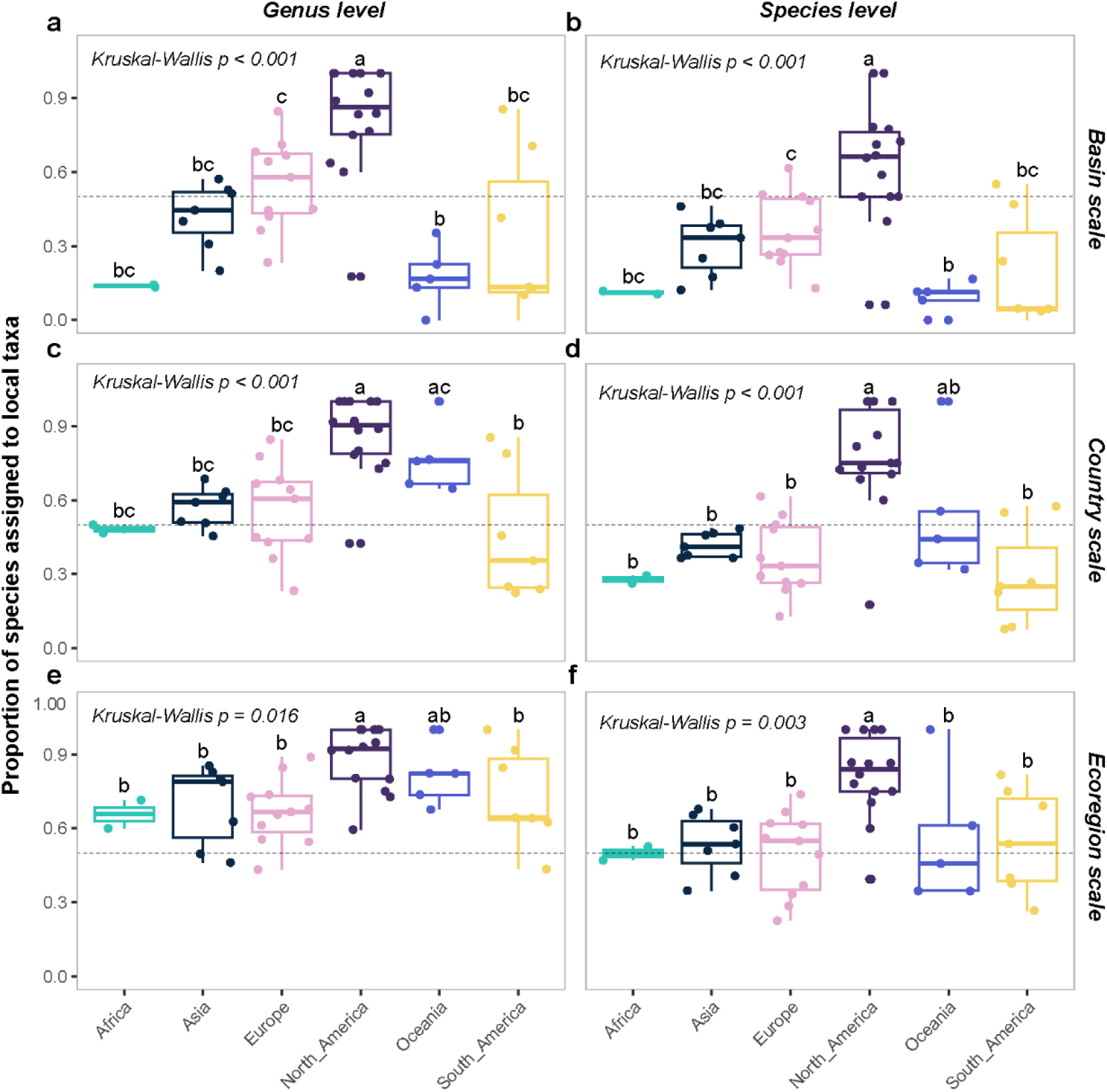
Proportion of species assigned as local taxa using global assignments. Local species were defined at the basin (a–b), country (c–d), and ecoregion (e–f) scales. Panels (a), (c), and (e) illustrate the genus-level comparisons, while panels (b), (d), and (f) show the species-level comparisons. The differences between ecoregions were assessed by ANOVA and the pairwise Dunn’s tests. Different letters indicate statistically significant differences between groups (p < 0.05). Groups sharing the same letter were not significantly different, whereas groups with no letters in common differed significantly. The horizontal dotted lines indicated a value of 0.5.

The validation of global species assignments using seven datasets with published historical fish records revealed notable differences in geographical scale and taxonomic resolution (Supplementary Table 4). At the basin scale, 74.5% to 100% of species-level assignments were validated as local. In contrast, the validation of genus-level assignments at the same scale exhibited greater variability. For the two datasets from Switzerland, 60% and 100% of assignments were validated as local genera, respectively. Contrastingly, datasets from French Guiana, St. Regis River, and Thailand showed much lower validation rates, ranging from 0% to 11.1%. Although historical records allowed for the validation of some species or genera at the country or ecoregion level, these taxa were not considered local at the basin scale, underscoring the necessity of basin-level data for accurate validation.

### Consistency in species richness estimates and community structure

The reanalyzed species richness estimates showed clear patterns across datasets but varied significantly depending on the spatial scales used to define local species. The original assignments identified a total of 1,521 species, with an average of 17 species per site (ranging from 0 to 106), while the reanalyzed global assignments identified 1,544 species, averaging 20 species per site (ranging from 0 to 337). When restricted to local species, the total species richness was reduced, with 895 species at the ecoregion scale (average 13 species per site, ranging from 0 to 190), 762 species at the country scale (average 11 species per site, ranging from 0 to 181), and 661 species at the basin scale (average 10 species per site, ranging from 0 to 149).

The LME results showed a high degree of consistency across all datasets and geographical scales, demonstrating the robustness of the species richness estimates (Figure 4, Supplementary Table 6). The global assignments, compared to the original assignments, showed strong performance (R^2^ = 0.22, R^2^ = 0.88, t = 20.64, *p* < 0.001). This consistency gradually decreased at finer scales, with assignments at the ecoregion scale maintaining relatively high explanatory power (R^2^ = 0.26, R^2^ = 0.86, t = 20.73, *p* < 0.001) and further declining at the country (R^2^ = 0.31, R^2^ = 0.82, t = 20.15, *p* < 0.001) and basin (R^2^ = 0.20, R^2^ = 0.84, t = 19.57, *p* < 0.001) scales.

**Figure 4.**
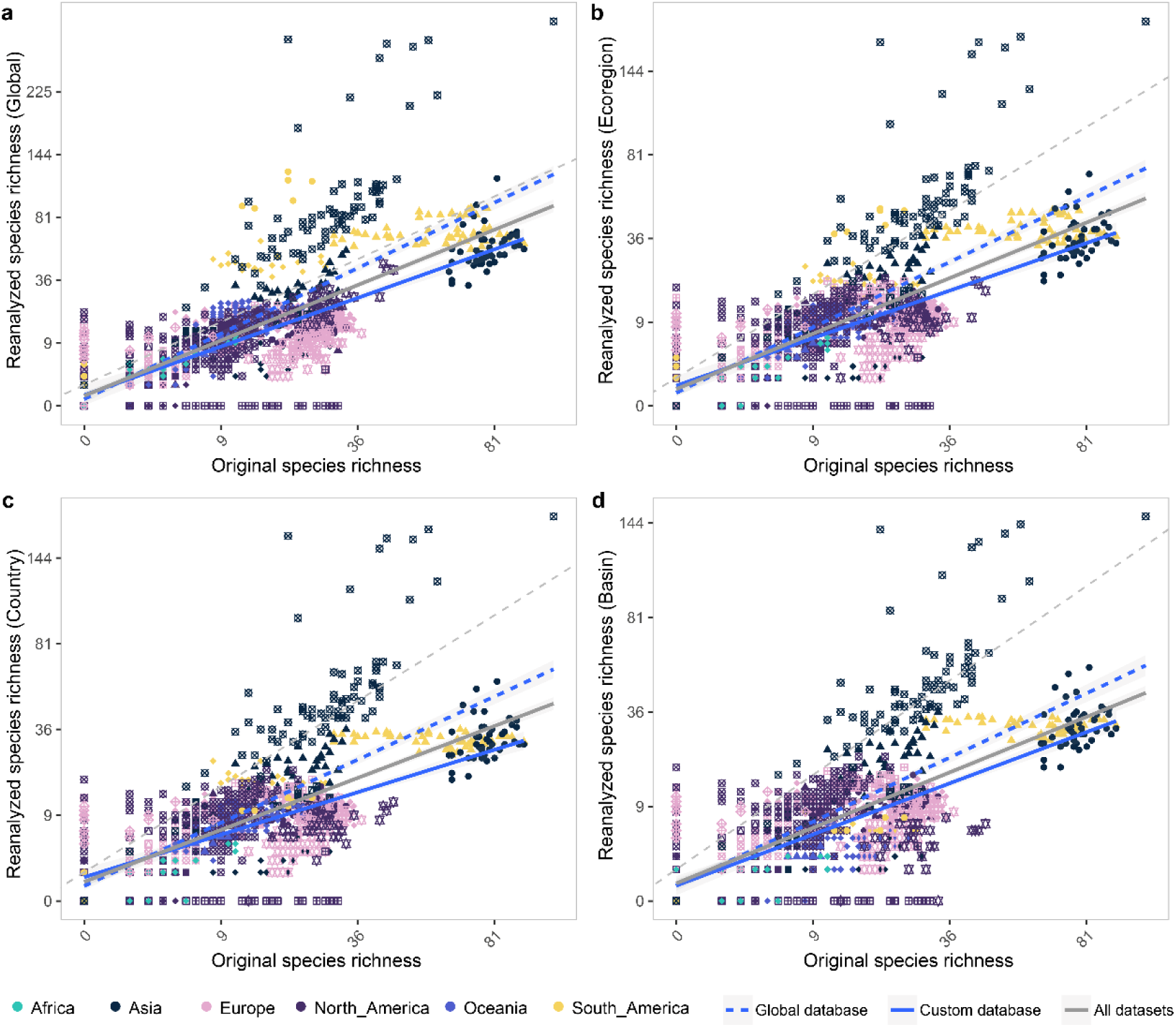
Comparison of species richness estimates between original and reanalyzed assignments. Panel (a) shows species richness based on global species assignments. Panels (b) to (d) display species richness estimates reanalyzed by restricting to local species at the ecoregion (b), country (c), and basin (d) scales. The grey dashed line represents 1:1 line. All regressions were performed using linear mixed models (LME), with dataset identity as a random factor to account for variation between datasets. The grey solid lines represent the regression for all datasets, the blue solid lines correspond to datasets that originally used custom reference databases for species assignment, and the dashed blue lines represent datasets originally using global reference databases. Point shapes corresponded to different dataset IDs for which raw sequence data were available.

When splitting the datasets based on their use of global or custom reference databases in the original assignments, notable differences emerged in the consistency of the species richness estimates (Figure 4, Supplementary Table 6). For datasets using the global reference database, the results showed strong agreement with the original assignments across all scales. At the global scale, the LME results for the global reference databases showed high performance (R^2^ = 0.59, R^2^ = 0.84, t = 22.67, *p* < 0.001), with consistency gradually decreasing at ecoregion (R^2^ = 0.56, R^2^ = 0.84, t = 22.29, *p* < 0.001), country (R^2^ = 0.54, R^2^ = 0.85, t_791.45_ = 21.96, *p* < 0.001), and basin scales (R^2^ = 0.49, R^2^ = 0.86, t_842.50_ = 21.51, *p* < 0.001). In contrast, datasets using custom reference databases showed less robust consistency, with t-values consistently below 10 and marginal *R*^2^ values explaining less than 1% of the variance (yet the conditional R^2^ above 0.90) at all scales. The weaker consistency for custom reference databases became more evident as the analysis shifted from global to local scales, with assignments deviating further below the 1:1 line.

Overall community composition remained consistently robust across species assignment types and was primarily structured by ecoregions (Figure 5, Supplementary Table 7). PERMANOVA results revealed a significant effect of ecoregions on community composition (R² = 0.19, *p* = 0.001), while species assignment types did not have a significant impact (R² = 0.019, *p* = 0.137). Similarly, ANOSIM confirmed a significant difference in community composition between ecoregions (R = 0.57, *p* = 0.001), but found no significant differences between species assignment types (R = 0.0031, *p* = 0.346).

**Figure 5.**
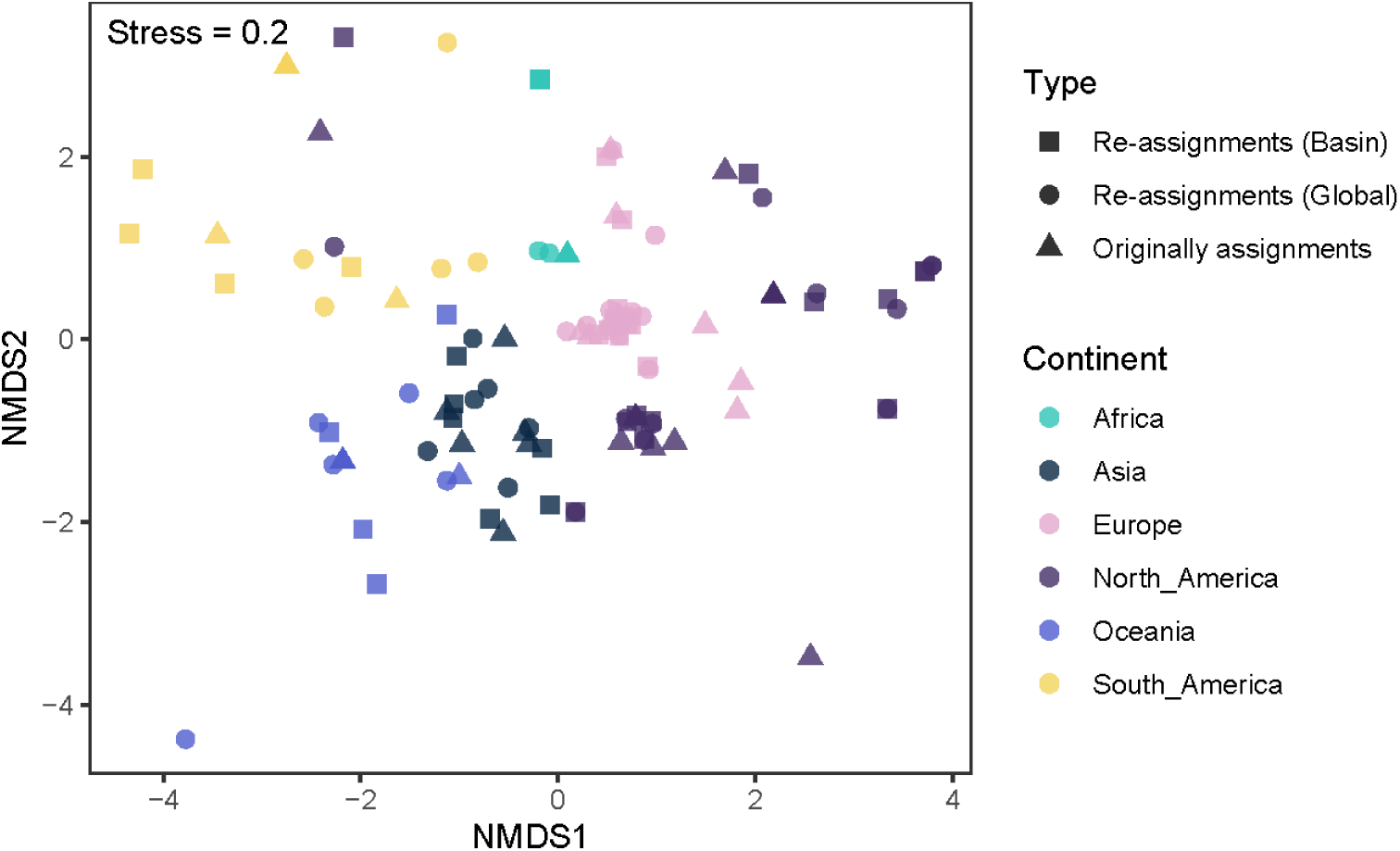
NMDS plots showing community composition across datasets. The colors represent different ecoregions, and the shapes indicate the dataset type: original assignment, global re-assignments, and basin-scale re-assignments.

### Comparison of species identity across eDNA studies

Out of the 50 reanalyzed datasets with raw sequencing data available reanalyzed, 46 were successfully annotated to basin-level local fish species. Among these 46 datasets, 40 had original species assignment tables available, enabling a direct comparison between the original and reanalyzed species identities (Supplementary Table 1). The species shared between the original and reanalyzed assignments ranged from 0 to 58 species per dataset (Figure 6 and Supplementary Figure 4), corresponding to 0–100% of species counts (mean: 60.71%, median: 66.67%). These shared species accounted for 0% to 100% of sequencing reads, with a mean and median overlap of 65.3% and 82.7%, respectively (Supplementary Figure 5). Significant differences were observed between ecoregions for the percentage of shared species in the reanalyzed assignments (Supplementary Table 8, Kruskall-Wallis H = 12.14 *p* = 0.033), while no significant differences were found between datasets original analyzed using global versus custom reference databases (Supplementary Table 9, Wilcoxon *p* = 0.734)

**Figure 6.**
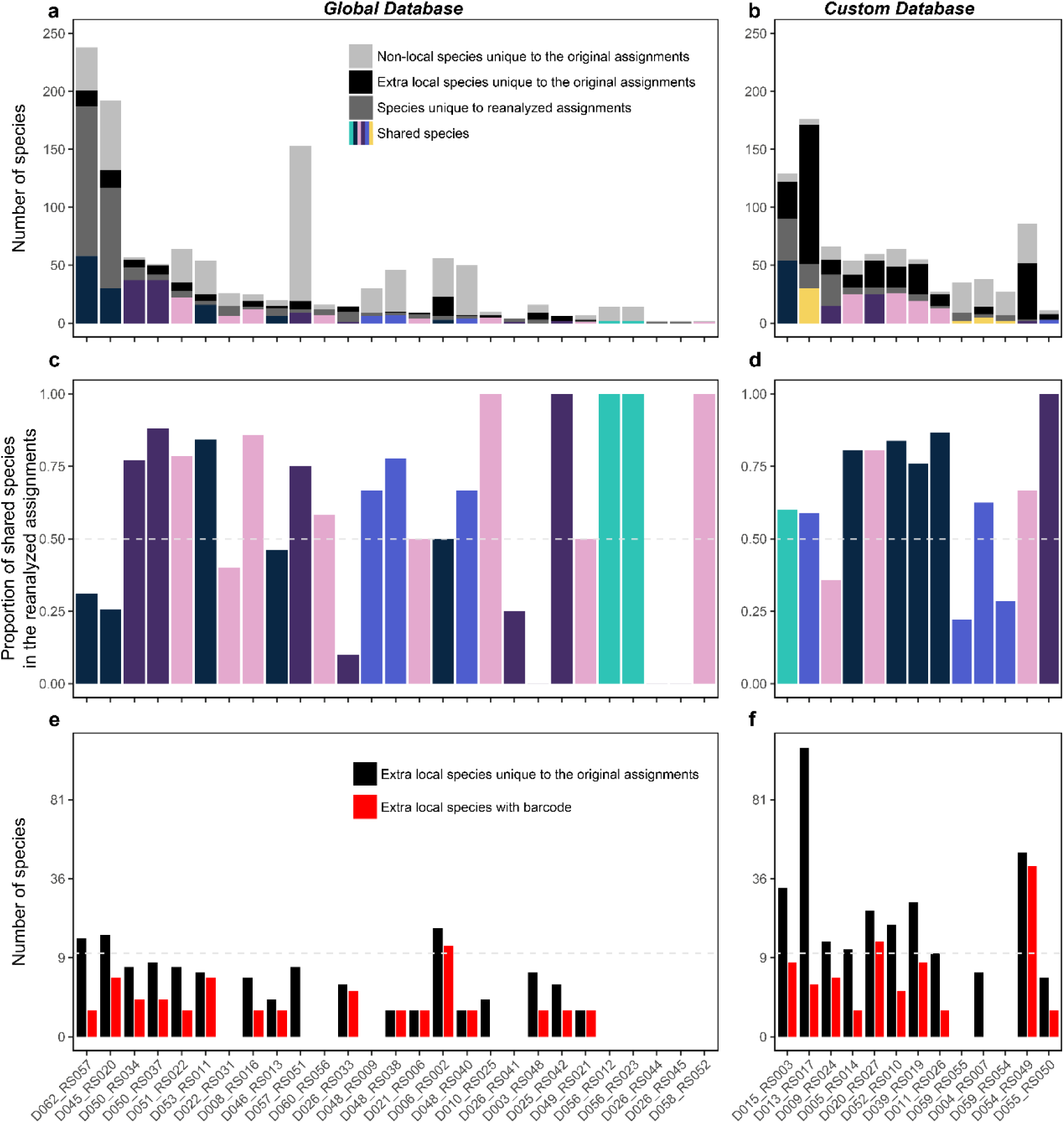
Comparison of species identity between the original and reanalyzed assignments for each dataset. The reanalyzed assignments only include species historically recorded at the basin scale. Panels (a–b) show the overlap between the original and reanalyzed assignments, with shared species color-coded by ecoregion. Dark grey bars represent species unique to the reanalyzed assignments, black bars indicate species unique to the original assignments but historically recorded at the basin scale (extra local species), and light grey bars represent species unique to the original assignments but not historically recorded. Panels (c–d) show the percentages of species detected in the reanalyzed dataset that were also found in the original dataset. Panels (e–f) show the number of extra local species in the original assignments, along with the number of these species with barcode records in GenBank. The x-axis in all panels represents individual dataset IDs. Assignments were classified based on the original assignment using either a global reference database (a, c, e) or a custom reference database (b, d, f).

The modelling results revealed that discrepancies between original and reanalyzed assignments were largely driven by sequencing depth, sampling year, and species richness, whereas the reference database type had minimal impact (Table 1, Supplementary Figure 6). The proportion of shared species (based on species count) increased significantly with higher median sequencing depth (t = 2.89, *p* = 0.007) and more recent year of sampling (t = 3.06, *p* = 0.004), but decreased with greater reanalyzed species richness (t = –2.91, *p* = 0.006). In contrast, the proportion of reads assigned to shared species was significantly affected only by the year of sampling, showing a higher overlap in more recent years (t = 2.74, *p* = 0.010). Reference database type (global vs. custom) had no significant effect in either model. Species unique to the reanalyzed datasets had significantly lower total relative abundance (z = –3.67, *p* < 0.001) and lower mean relative abundance across sampling sites (z = –4.07, *p* < 0.001) compared to shared species (Supplementary Figure 7 and Supplementary Table 10). Reference database type showed marginal effects on total (*p* = 0.083) and mean (*p* = 0.055) relative abundance, whereas the year of sampling had no significant effect.

**Table 1.**
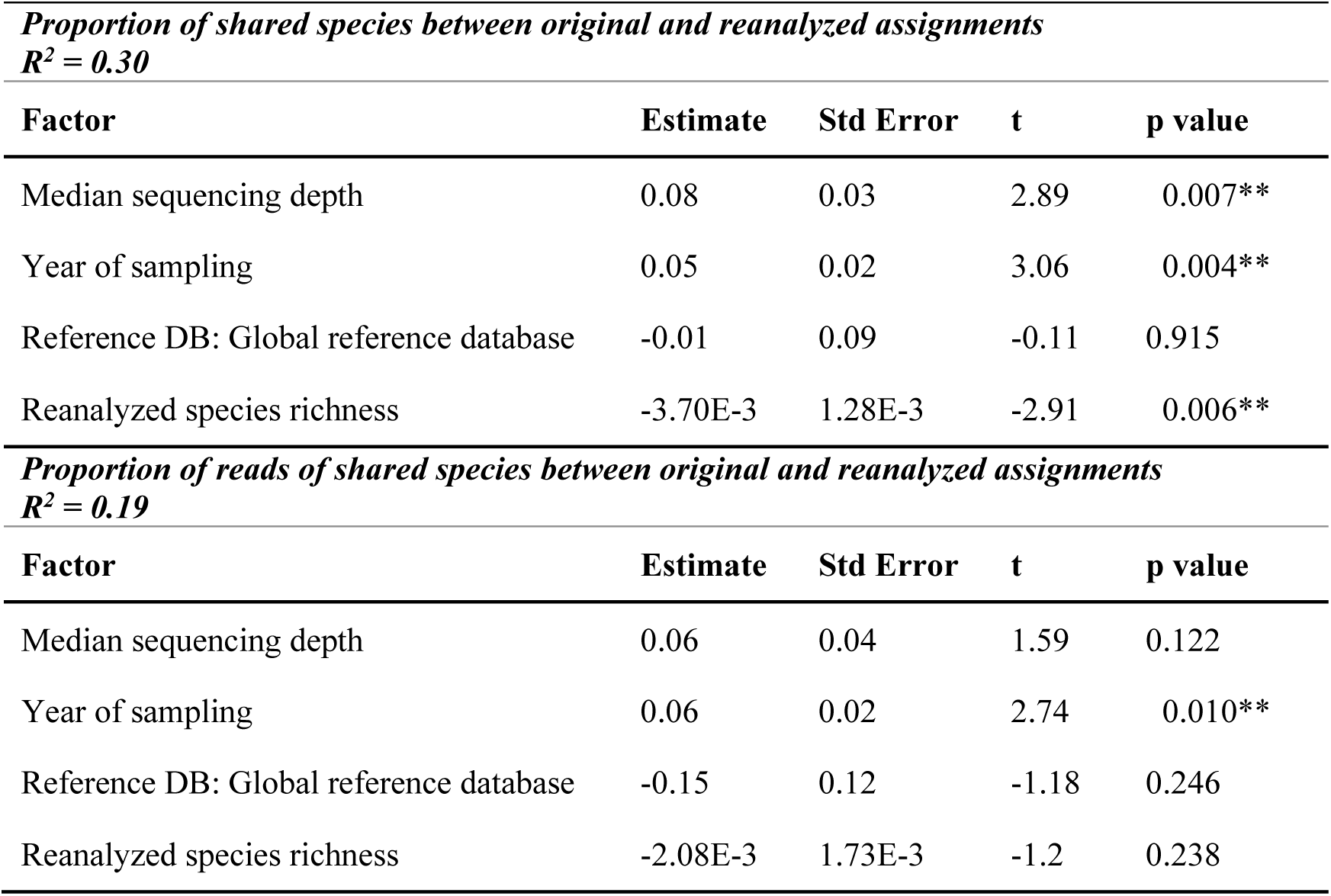
Results of generalized linear models (GLMs) testing factors influencing the proportion of shared species between original and reanalyzed assignments, based on (a) species count and (b) sequencing read abundance.

Despite the overlap, some species were unique to the original assignments as basin-scale local species (extra local species hereafter), with 11 species per dataset on average, ranging from 0 to 120 (Figure 6, Supplementary Figure 4). No significant differences in the number of extra local species were found between ecoregions (Supplementary Table 8, Kruskall-Wallis H = 15.94 *p* = 0.159), but datasets using custom reference databases in the original assignments had significantly more extra local species (range from 0 to 120) compared to those using global reference databases (Supplementary Table 9, range from 0 to 17; Wilcoxon *p* = 0.003). In general, only 0 to 13 of these extra local species appeared in the curated reference database used for reanalysis. An exception was dataset D054, which had the lowest reads per sample (2,597 reads, Supplementary Table 11), resulting in 4 out of 443 ASVs being identified as fish (Supplementary Figure 8). In contrast, other datasets had an average of 9,943 to 3,704,035 reads per sample (Supplementary Table 11).

## Discussion

Environmental DNA has become a powerful tool for monitoring riverine fish biodiversity, providing a non-invasive approach for large-scale assessments (Blackman et al., 2024). However, questions remain about its global applicability, particularly the consistency of existing datasets generated through varying bioinformatic workflows. The storage and availability of raw sequence data have been critical in enabling this reanalysis, and our study stands as one of the first and largest efforts to reanalyze fish eDNA datasets under a common bioinformatic framework. Specifically, our major reanalysis revealed that overall biodiversity patterns in terms of species richness and community structure were consistent, yet discrepancies did arise in the species identities between the reanalyzed and original assignments. These differences were not necessarily due to variations in the bioinformatic pipeline, but rather stemmed from the choice and quality of reference databases. These findings underscore the potential of eDNA as a reliable and scalable tool for monitoring global fish biodiversity, while highlighting the importance of refining bioinformatic workflows and reference database curation for more accurate species identification.

A key finding of this study is that eDNA-derived biodiversity patterns were generally robust, underscoring the potential of leveraging existing fish eDNA datasets for global biodiversity analyses. As eDNA metabarcoding continues to advance in biodiversity monitoring, concerns linger about how methodological factors, including bioinformatic approaches, may influence local diversity estimates (Blackman et al., 2024; Marques et al., 2020). Various bioinformatic tools, such as MiFish pipeline (Sato et al., 2018), OBITools (Boyer et al., 2016), Anacapa (Curd et al., 2019), and QIIME (Bolyen et al., 2019; Caporaso et al., 2010), offer comprehensive eDNA analysis workflows with the same primary objectives—inferring species identities and estimating species richness— but they employ different strategies (Hakimzadeh et al., 2024; Mathon et al., 2021). These workflows diverge in several critical areas, including OTU/ASV clustering methods, taxonomic annotation strategies, and criteria for defining taxonomic resolution (Hakimzadeh et al., 2024; Mathon et al., 2021). Moreover, the choice of reference databases is crucial for eDNA metabarcoding analysis (Keck & Altermatt, 2023; Keck et al., 2022b). Public reference databases offer greater taxonomic breadth but tend to have lower resolution for shorter eDNA barcodes and higher rates of errors/ taxonomic misidentifications, which can result in misidentifications or false positives (Keck et al., 2022b). On the other hand, local reference databases, though narrower in scope, tend to provide more accurate species identification due to their focus on regional taxa. Given these differences, the impact of bioinformatic workflows on the accuracy and consistency of biodiversity assessments is one of the central concerns when compiling eDNA datasets for global biodiversity insights (Blackman et al., 2024; Hakimzadeh et al., 2024; Mathon et al., 2021). Our reanalysis revealed the consistency of species richness estimates and the robustness of community structures across studies employing unified bioinformatic workflow (Figure 4 and Figure ***5***), indicating existing eDNA data can offer reliable community-level insights.

The choice and quality of reference databases critically influenced the accuracy of species identification in eDNA studies, with ongoing improvements over time helping to reduce taxonomic mismatches (Keck et al., 2022b). In our reanalyses, species identity mismatches between original and reanalyzed datasets varied across datasets, largely reflecting differences in how reference databases were constructed and applied (Figure 6). First, our reanalyses revealed that restricting species identification to basin-scale local species pools substantially improved taxonomic resolution, emphasizing the value of regionally tailored reference databases for accurate biodiversity assessments. For instance, datasets originally analyzed using global reference databases often showed improved species annotations when reanalyzed with basin-specific species lists, underscoring the importance of aligning reference data with local faunas. Conversely, in regions such as South America, where custom reference databases were used but not publicly archived (e.g., Cantera et al. (2022)), reanalyses suffered from lower taxonomic resolution due to limited access to these localized resources. Second, persistent gaps and geographic biases in reference database coverage remained a significant challenge. Previous studies have highlighted that gaps in reference database coverage for fish species were especially prominent in tropical regions, where both species diversity and the number of threatened species were highest (Marques et al., 2021). In Europe, while freshwater fish species were relatively well represented with 88% coverage of COI gene, the coverage for other markers such as 12S rRNA remains low, with only 36% of species covered (Weigand et al., 2019). Similarly, for Chinese fish species, around 60% were represented in reference databases, but nearly 90% of the sequences came from outside China, highlighting a critical geographical bias (Li et al., 2022). Our study found the uneven coverage of reference databases across ecoregions, with notable gaps in the taxonomic representation of local fish species in South America and Africa, was the main reason for the species identities differences between original and reanalyzed results (Figure 2). Third, our results showed that the year of sampling was positively associated with species identification accuracy, indicating that improvements in reference databases over time could enhance biodiversity estimates (Table 1). Reanalysis of “old” datasets with updated reference libraries thus provides a valuable opportunity to improve consistency and comparability across studies. Overall, our findings highlighted the need for continued expansion and curation of region-specific reference databases to support reliable and standardized eDNA-based biodiversity monitoring.

While eDNA data provides a promising tool for global biodiversity assessments, the growing issue of spatial bias in eDNA research raises concerns. In the context of global biodiversity targets, such data discrepancies could lead to the systematic underestimation of biodiversity in underrepresented regions, thereby introducing bias into global biodiversity evaluations and subsequent scientific decision-making (Chapman et al., 2024). Traditional biodiversity monitoring data, as represented by the Global Biodiversity Information Facility (GBIF), has already demonstrated geographic bias, with most data originating from high-income regions (Chapman et al., 2024; Hughes et al., 2021a). Therefore, in addition to the need for more biodiversity data, there is an urgent need to focus on underrepresented areas and taxa. Our meta-analysis shows that, aside from the geographic bias in the reference databases mentioned earlier, the geographic distribution of eDNA datasets also exhibits spatial bias, with a concentration of studies in Europe, North America, and East Asia, while regions such as South America, Africa, Oceania, and other parts of Asia remain under-sampled (Figure 1). Similar to the study by Keck et al. (2022a), eDNA metabarcoding studies, though covering multiple continents and climatic regions, still show a marked spatial clustering, with most studies focused on Europe and North America. Research on airborne bacterial eDNA also reveals a geographical bias, with more data from Europe and Asia, fewer from North America, and almost no data from Africa, South America, and Oceania (Jiang et al., 2022). This highlights the critical need for future eDNA research to address these geographic disparities by increasing efforts in underrepresented regions. Doing so will be essential to support a more comprehensive, equitable, and accurate global biodiversity conservation agenda.

Our reanalysis underscored the critical importance of raw data availability and comprehensive metadata in future eDNA studies. Not all datasets could be effectively used in the reanalysis. Out of the 42 published datasets examined, raw sequence data were missing in 17 cases, while an additional four datasets included raw sequencing data but lacked barcodes/indexes necessary to distinguish individual samples from sequencing files. Moreover, the absence of key metadata in some datasets hindered the reanalysis process. For example, some studies employed modified primer sets but failed to report the exact primer sequences, instead referencing the original publication. Other issues included inconsistencies between sequencing file names and specific samples, as well as the omission of sampling site coordinates. This highlights a critical issue: for eDNA data to be truly useful, careful management and appropriate data storage protocols are essential (Altermatt et al., 2025). In this study, we outlined key metadata including sampling, metabarcoding experiment process and detailed bioinformatics pipeline that should be reported to facilitate and enhance future reanalysis (Supplementary Table 1). Looking ahead, it is crucial to establish standardized data storage protocols and ensure the reporting of comprehensive metadata for reanalysis (see Klymus et al. (2024)), similar to the practices already implemented in platforms such as GBIF (Telenius, 2011), eBird (Tang et al., 2021), and NCBI (Jenuth, 1999). By adopting such measures, eDNA can truly deliver valuable data and insights, contributing meaningfully to global biodiversity monitoring and assessment efforts (Altermatt et al., 2025; Thomsen et al., 2024).

## Acknowledgments

This study is supported by the National Key Research and Development Program of China (2022YFC32021001, 2021YFC3201003) and National Natural Science Foundation of China (42507382). We thank all the many people involved in the field and laboratory work of the respective studies we base our reanalysis on. We also thank all researchers who provided comments and information on the published dataset. Funding is from the Swiss National Science Foundation, grants 310030_197410 and 31003A_173074 (to FA). The views expressed in this article are those of the author(s) and do not necessarily represent the views or policies of the U.S. Environmental Protection Agency.

## Author Contributions

Y.Z. and F.A. conceived and led the study. Y.Z. collected and analyzed the datasets, and wrote a first version of the manuscript, with input and mentoring of F.A. and X.Z. H.Z. assisted with data cleaning. All other authors contributed data. All authors commented on drafts of the paper.

